# Genetic Deletion of Skeletal Muscle Inositol Polyphosphate Multikinase Disrupts Glucose Homeostasis and Impairs Exercise Tolerance

**DOI:** 10.1101/2024.07.28.605526

**Authors:** Ji-Hyun Lee, Ik-Rak Jung, Becky Tu-Sekine, Sunghee Jin, Frederick Anokye-Danso, Rexford S. Ahima, Sangwon F. Kim

**Affiliations:** Department of Medicine, Division of Endocrinology, Diabetes, and Metabolism, Johns Hopkins University, Baltimore, Maryland, USA. 21224

**Author notes:** **Contact Information:** Corresponding Authors: Sangwon F. Kim, PhD and Rexford S. Ahima, MD, PhD Department of Medicine, Division of Endocrinology, Diabetes, and Metabolism, Johns Hopkins University, 5501 Hopkins Bayview Cir, AAC 2A44, Baltimore, Maryland, USA. 21224, and.

**Keywords:** Inositol polyphosphate, IPMK, skeletal muscle, exercise, insulin

## Abstract

Inositol phosphates are critical signaling messengers involved in a wide range of biological pathways in which inositol polyphosphate multikinase (IPMK) functions as a rate-limiting enzyme for inositol polyphosphate metabolism. IPMK has been implicated in cellular metabolism, but its function at the systemic level is still poorly understood. Since skeletal muscle is a major contributor to energy homeostasis, we have developed a mouse model in which skeletal muscle IPMK is specifically deleted and examined how a loss of IPMK affects whole-body metabolism. Here, we report that mice in which IPMK knockout is deleted, specifically in the skeletal muscle, displayed an increased body weight, disrupted glucose tolerance, and reduced exercise tolerance under the normal diet. Moreover, these changes were associated with an increased accumulation of triglyceride in skeletal muscle. Furthermore, we have confirmed that a loss of IPMK led to reduced beta-oxidation, increased triglyceride accumulation, and impaired insulin response in IPMK-deficient muscle cells. Thus, our results suggest that IPMK mediates the whole-body metabolism via regulating muscle metabolism and may be potentially targeted for the treatment of metabolic syndromes.

## INTRODUCTION

Inositol Polyphosphate Multikinase (IPMK) is the rate-limiting enzyme in the synthesis of soluble inositol polyphosphates (IPs) and inositol pyrophosphates (PP-InsPs), and an important component in the recycling of phosphatidylinositol lipids (PIPs) [1–6]. Targeted disruption of various enzymes that participate in IP synthesis has shown that these molecules impact a variety of cellular processes ranging from gene transcription to cellular metabolism, and IPs are now fully recognized as critical second messengers [7–12]. IPK2, the yeast homolog of mammalian IPMK, was originally identified as a factor (Arg82) necessary for the assembly of the ArgR/Mcm1 complex, which reciprocally regulates the expression of anabolic and catabolic genes required for arginine metabolism, exerting both scaffolding and activity-dependent functions [13, 14]. Murine IPMK also participates in metabolic functions, including amino acid sensing and glucose homeostasis through protein-protein interactions (PPIs) with AMPK, LKB1, and mTORC1. In addition, IPMK plays a critical function in regulating growth factor signaling, including insulin response [7, 15–18].

Increased skeletal muscle mass and physical activity are beneficial for metabolic, cardiovascular, and overall health [15–17]. Adult skeletal muscles are composed of a variety of fiber types, each specialized in terms of morphology, contractile properties, metabolism, and resistance to fatigue, enabling them to undertake specialized functions [18, 19]. Skeletal muscle comprises ∼40% of body mass in mammals, is the major determinant of energy expenditure at rest and during physical activity, and plays a critical role in glucose homeostasis as the predominant site of insulin-stimulated glucose disposal and glycogen storage [20, 21]. Skeletal muscle contributes ∼30% of whole-body metabolism at rest and up to ∼90% during maximal exercise [22].

Various signaling pathways have been linked to skeletal muscle function [23–26]. AMPK is a serine/threonine that modulates cellular metabolism acutely via phosphorylation of metabolic enzymes and long-term via transcriptional regulation [27–30]. AMPK is activated by cellular energy deficit, e.g., fasting and exercise, resulting in an increased AMP/ATP ratio. Acute AMPK activation suppresses glycogen and protein synthesis, and promotes glucose and fatty acid oxidation, while chronic AMPK activation induces mitochondrial biogenesis through co-activator PGC-1α [31, 32] and alters metabolic gene expression via transcription factors[27, 28]. Given the interactions of IPMK with key metabolic regulators, i.e., Akt, AMPK, and mTOR [4, 7, 33, 34], we wondered whether skeletal muscle IPMK may play a critical role in whole-body metabolism. Here, we report that a loss of skeletal muscle IPMK (henceforth referred to as MKO) disrupts insulin sensitivity and lipid utilization, leading to an increase in body weight, glucose intolerance, and reduced exercise tolerance. Thus, we elucidate the role of skeletal muscle IPMK in muscle energy metabolism through the regulation of insulin response and lipid utilization.

## Methods and Materials

### Mice

All animal experimental procedures were approved by the Johns Hopkins University Animal Care and Use Committee. To generate adipocyte-specific *Ipmk* knockout mice, *Ipmk*-floxed mice were mated with MLC-Cre transgenic mice (a generous gift from Dr. Se-Jin Lee, University of Connecticut) (MKO). *Ipmk-*floxed mice were developed, as described previously [35] and backcrossed with C57BL/6J wild-type for at least five generations. All mice were maintained in a 12:12-h light-dark cycle with free access to regular chow (4.5% fat, 49.9% carbohydrate, 23.4% protein; 4 kcal/g, LabDiet, Richmond, IN) and water in a specific pathogen-free facility in the Johns Hopkins University.

### Primary myocytes

Skeletal muscle tissues were isolated from 3-5 neonate (1-3 days old) pups and minced with a blade. Tissues were mixed with add 2ml Collagenase/Dipase/CaCl_2_ solution per gram and incubated at 37°C for 20 mins. Myocytes were collected by centrifugation at 350g for 5 min and resuspended in F-10 based primary myoblast growth medium (Ham’s F-10 nutrient mixture, 20% fetal calf serum, 0.5% bovine serum, 1% penicillin/streptomycin (100U/ml &100ug/ml each) in a tissue culture dish coated with collagen. Incubate in 37°C 5% CO_2_ incubator and change medium every 2 days. After fibroblasts are no longer visible, the medium was switched to F-10/DMEM based primary myoblast growth medium (F-10:DMEM=1:1) daily. For differentiation, medium was replaced with Fusion Medium (DMEM with 5% horse serum and penicillin/streptomycin).

### Measurement of body composition, energy balance and Indirect colorimetry

Body composition was measured by EcoMRI (dual-energy X-ray absorptiometry) scan. Energy expenditure was measured by open-circuit calorimetry (Oxymax system, Columbus Instruments, Columbus OH) and locomotor activity was measured simultaneously by infrared beam interruption (Opto-Varimex System, Columbus Instruments, OH). Mice were housed individually in calorimetry cages at 23°C. Room air was pumped at a rate of 0.52 liters/min and exhaust air was sampled at 15 mins intervals for 5 hours. Oxygen consumption (VO_2_) and carbon dioxide production (VCO_2_) were measured using electrochemical and spectrophotometric sensors, respectively. Respiratory quotient (RQ), a measure of fuel use, was calculated as the ratio of oxygen consumption to carbon dioxide production. A decrease in RQ indicates fatty acid oxidation. Total energy expenditure (heat) = Calorific value (CV) × VO_2_, where CV = 3.815 + 1.232 × RQ.

### Treadmill exercise

For assessment of muscle endurance capacity, WT and IPMK MKO male mice (n = 8 per group) were tested for treadmill running. For the treadmill running, an Open Rodent Treadmill Exer-3/6 was used. Prior to exercise, mice were accustomed to the treadmill with a 2-3 min explore once per day for 3 days. The exercise test was performed on a 10% incline for 10 m/min for 5mins, followed by an increase toward 30 m/min until exhaustion.

### Glucose tolerance test

Mice were fasted for 6 hours and glucose (2 g/kg body weight; i.p.) was administered intraperitoneally. After injection, blood was taken by puncturing of the tail vein, and glucose levels were measured using a glucometer (Ascensia Diabetes Care) at 0, 15, 30, 60 and 120 mins. Blood glucose was also measured before the injection (time point 0).

### Quantitative Real time-PCR

Total RNAs were isolated using TRIzol Reagents. RNAs were reverse transcribed into cDNA using Protoscript II (New England BioLab). 1-2 μg of cDNA was diluted to 2 ng/ml and was amplified by specific primers in a 10 µl reaction using SYBR Green Master mix (Applied Biosystems). Analysis of gene expression was carried out in QuantStudio 5 (Applied Biosystems). Mouse TATA-binding protein gene (*Tbp*) was used as the reference gene, and data was normalized and relative expression determined from the threshold cycle (Ct) following the 2 ^−ΔΔCT^ method. Primers were as follows: SCD1 forward 5’-GCAAGCTCTACACCTGCCTCT-3’, reverse 5’-CGTGCCTTGTAAGTTCTGTGGC-3’; DGAT1 forward 5’ - TGACCTCAGCCTTCTTCCATGAGT-3’, reverse 5’-CCACACAGCTGCATTGCCATAGTT-3’; CPT1b forward 5’-GGCACCTCTTCTGCCTTTAC-3’, reverse 5’-TTTGGGTCAAACATGCAGAT-3’; CD36 forward 5’-ACTGGTGGATGGTTTCCTAGCCTT-3’, reverse 5’-TTTCTCGCCAACTCCCAGGTACAA-3’; FATP1 forward 5’-CCGTCTGGTCAAGGTCAATG-3’, reverse 5’-CACTAACATAACCATCGAAACGC-3’.

### Western blotting

Samples were lysed in ice-cold RIPA buffer containing protease and phosphatase inhibitors and heated at 95°C for 5 min prior to electrophoresis. Proteins were transferred to a 0.2 µm nitrocellulose membrane, blocked with 5% nonfat milk in Tris-buffered saline for 30 mins. Blots were then incubated with primary antibodies at 4°C. Immunoblotting was conducted with the following antibodies: IPMK from Novus Biologicals and pAMPK, AMPK, pAkt-S473, Akt, nSREBP, Myogenin, GAPDH from Cell Signaling Technology. Blots were imaged and quantitated using an Odyssey Near-Infrared Scanner (Li-Cor Biosciences).

### Glucose Uptake Assay

Cells were washed with PBS and incubated in serum free medium for 2 hrs. Then cells were incubated in Krebs-Ringer biocarbonate-Hepes buffer (KRBH, 30 mM/pH7.4 HEPES, 10 mM NaHCO_3_, 120 mM NaCl, 4 mM KH_2_PO_4_, 1 mM MgSO_2_, 1 mM CaCl_2_) with 1 mM 2-deoxyglucse plus 1μCi [14C]-2-deoxyglucose for 10 mins. Cells were washed with cold PBS (x3) and radioactivity was measured by scintillation counter. Glucose uptake was normalized by the protein concentration in each well.

### Fatty acid Uptake and Oxidation

The measurement of fatty acid oxidation was performed as previously described. Briefly, primary myocytes were starved for 2 hrs and then incubated for 4 hrs in presence of 5 mM glucose, 0.1 mM oleic acid pre-complexed to 0.13% BSA [1–14 C]-Oleic acid (0.1 μCi/ml). The trapped ^14^CO_2_ and 14C-Acid Soluble Products were counted to estimate total oleic acid oxidation using the liquid scintillation analyzer Tri-Carb 4810 TR. The results were normalized by cell protein content.

### Statistical Analysis

Results were analyzed using a statistics software package (GraphPad Prism version 9.0.0 for Windows, GraphPad Software, San Diego, California USA). We assessed statistical significance by unpaired two-tailed t-tests, one-way ANOVA (Dunnet’s Test), or two-way ANOVA (Tukey’s test) for multiple comparisons. The level of significance was set to p < 0.05.

## Results

### A loss of IPMK in skeletal muscle disrupts energy homeostasis

To better understand the metabolic role of IPMK, we have created skeletal muscle-specific Ipmk null mice by crossing *Ipmk-*loxp (neomycin deleted) mice with MLC-Cre. MKO mice were born without any obvious gross abnormality but started to gain more weight at 13-14 weeks (**Fig. 1A**). To assess energy homeostasis, MKO and WT mice were housed singly in metabolic cages. The respiratory exchange ratio (RER) in MKO was significantly elevated especially in the light period compared to WT, indicating impairment of fat oxidation (**Fig. 1B**). Spontaneous locomotor activity measured using photobeam breaks was not different between MKO and WT (**Fig. 1C**). Energy expenditure against lean mass was decreased in MKO mice (**Fig. 1D**). Food intake was slightly increased without any statistical significance in MKO (data not shown). These mice were further examined for body composition at 22 weeks. We found that the MKO gained more weight and exhibited an increased fat mass compared to WT. The lean mass in MKO mice was slightly decreased without statistical significance (**Fig. 1E, F**). Overall, these results indicated that MKO mice had reduced oxidative capacity with lower energy availability ultimately leading to weight gain.

**Figure 1.**
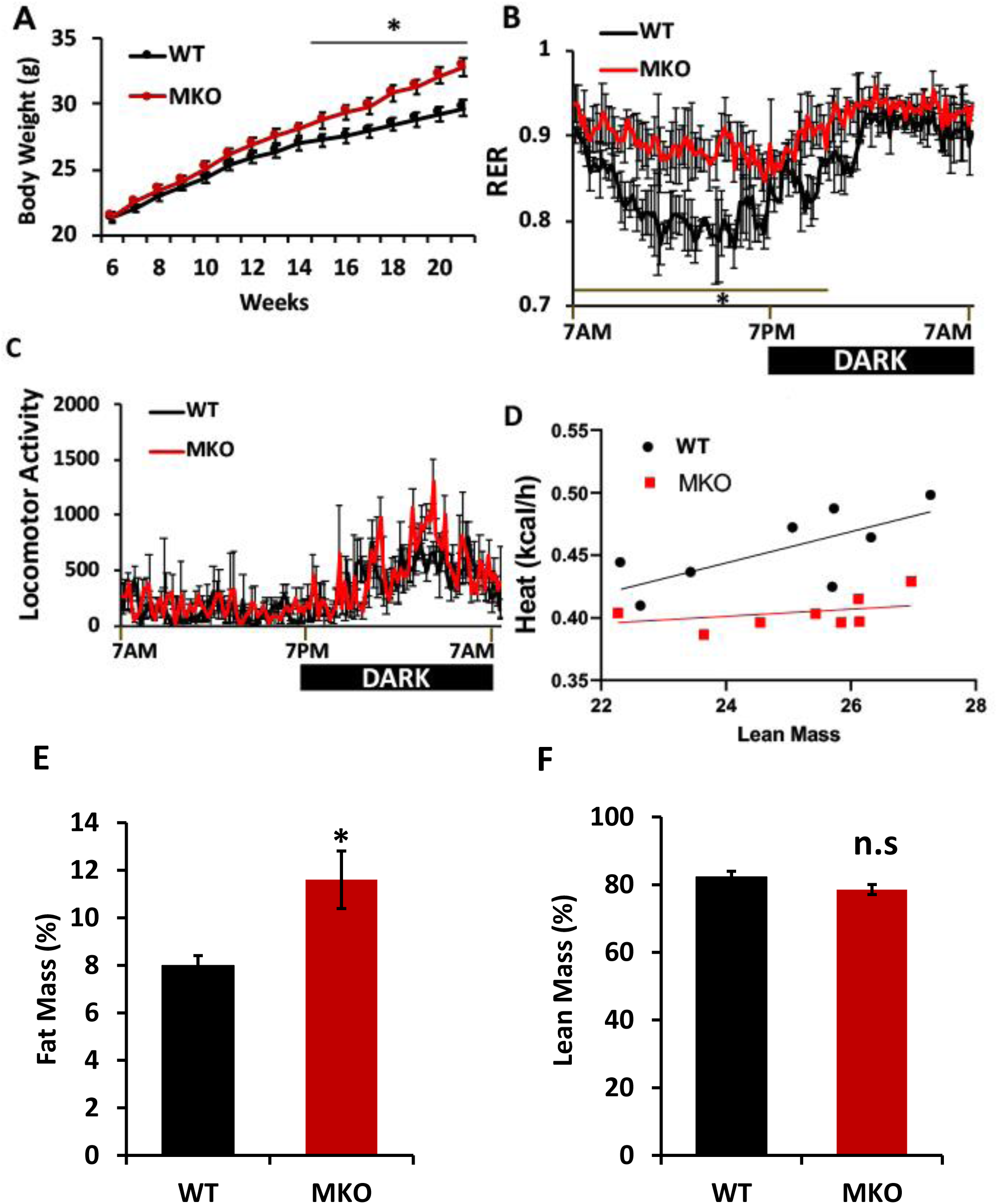
Effects of skeletal muscle IPMK deficiency in the whole-body energy metabolism. Male (22-24-week-old) WT and MKO mice were examined **A.** Body weight, **B.** Indirect calorimetry was performed respiratory exchange ratio (RER; VCO_2_/VO_2_) **C.** Locomotor activity measured with photobeam breaks. **D.** Regression analysis of energy expenditure **E and F.** Lean and fat mass were determined by ^1^H-MRS analysis. Comparisons between the two groups using Student’s t-test. The differences between genotypes over 24 h were analyzed by two-way ANOVA and Bonferroni post hoc test. Data are mean +/-SEM, (WT, n = 8; MKO, n = 8). *p<0.05.

### Glucose homeostasis is impaired in MKOs

Glucose is one of the main energy sources for the skeletal muscle through glycolysis and oxidative phosphorylation, and impaired glucose metabolism in muscles leads to reduced energy usage and increased fat storage, contributing to weight gain and insulin resistance [16, 36]. To study the effect of IPMK deletion in muscle on glucose homeostasis, we performed an intraperitoneal glucose tolerance test at 22 weeks of age. After 6 h of fasting, blood glucose levels in MKO were significantly higher than in WT (**Fig. 2A)**. Moreover, we observed that MKO developed markedly impaired glucose tolerance as compared to WT, suggesting that a loss of IPMK in skeletal muscle disrupts whole-body glucose metabolism (**Fig. 2B**).

**Figure 2.**
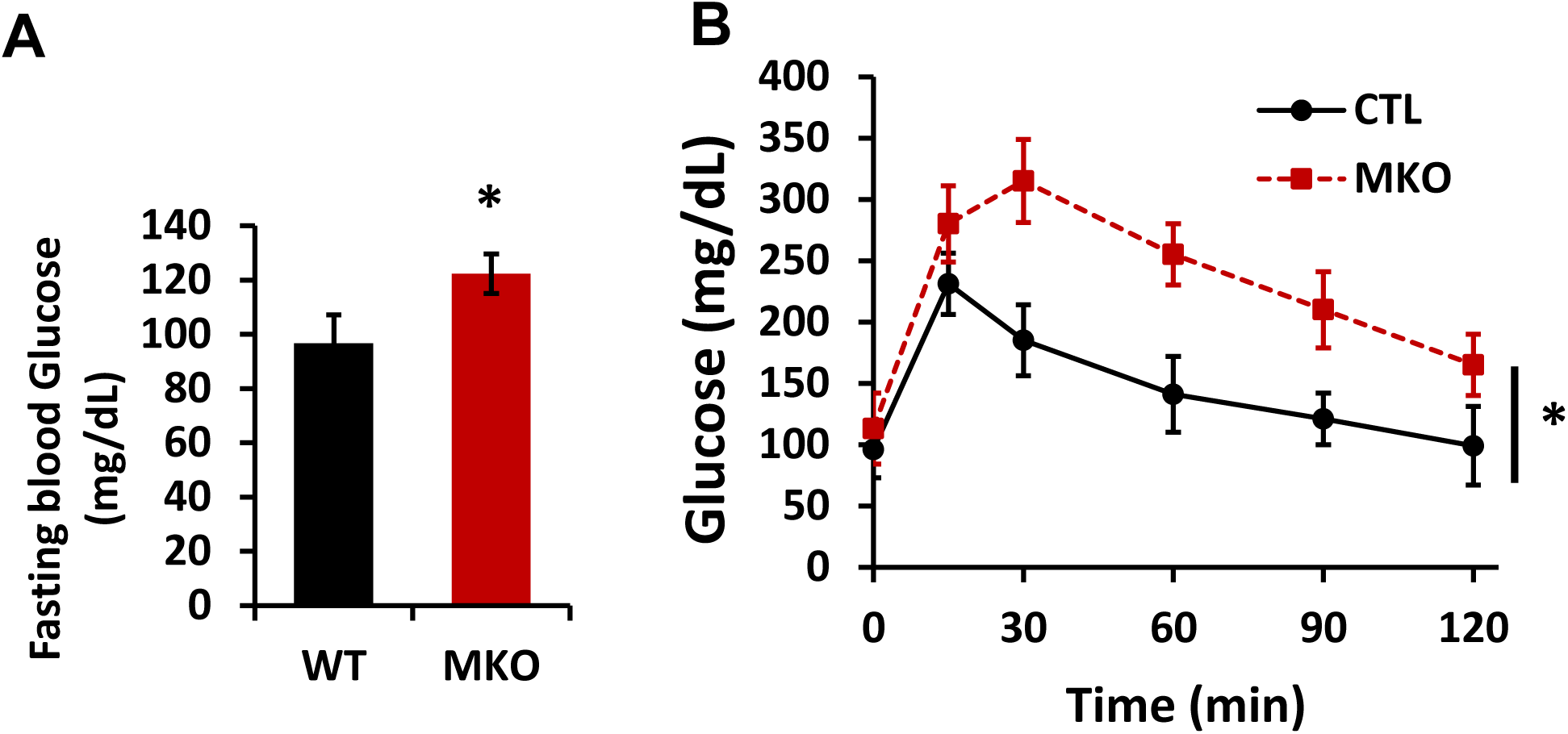
Effects of skeletal muscle IPMK deficiency on glucose homeostasis. Male (22-24-week-old) WT and MKO mice were examined. A. overnight fasting glucose and B. glucose tolerance test (GTT). Data are mean +/- SEM, (WT, n = 8; MKO, n = 8). *p<0.05.

### Deletion of skeletal muscle IPMK disrupts gene expression associated with lipid metabolism

Skeletal muscle also utilizes lipids as a critical energy source via beta-oxidation. Disrupted lipid metabolism in skeletal muscle leads to fat accumulation, insulin resistance, impaired energy production, and inflammation, contributing to metabolic disorders like obesity and type 2 diabetes [16, 37]. Hence, we examined the expression of genes related to lipid metabolism in gastrocnemius muscle. We noted that gene expression for lipid uptake (CD36) and synthesis (SCD1) was significantly increased, while the fat oxidation gene (CPT1b) was decreased in MKO mice **(Fig. 3A).** These changes were further associated with an increase in triglyceride (TG) in the muscle tissue in MKO **(Fig. 3C).** These data suggest that a loss of skeletal muscle IPMK disrupts how muscle utilizes the lipid as an energy source.

**Figure 3.**
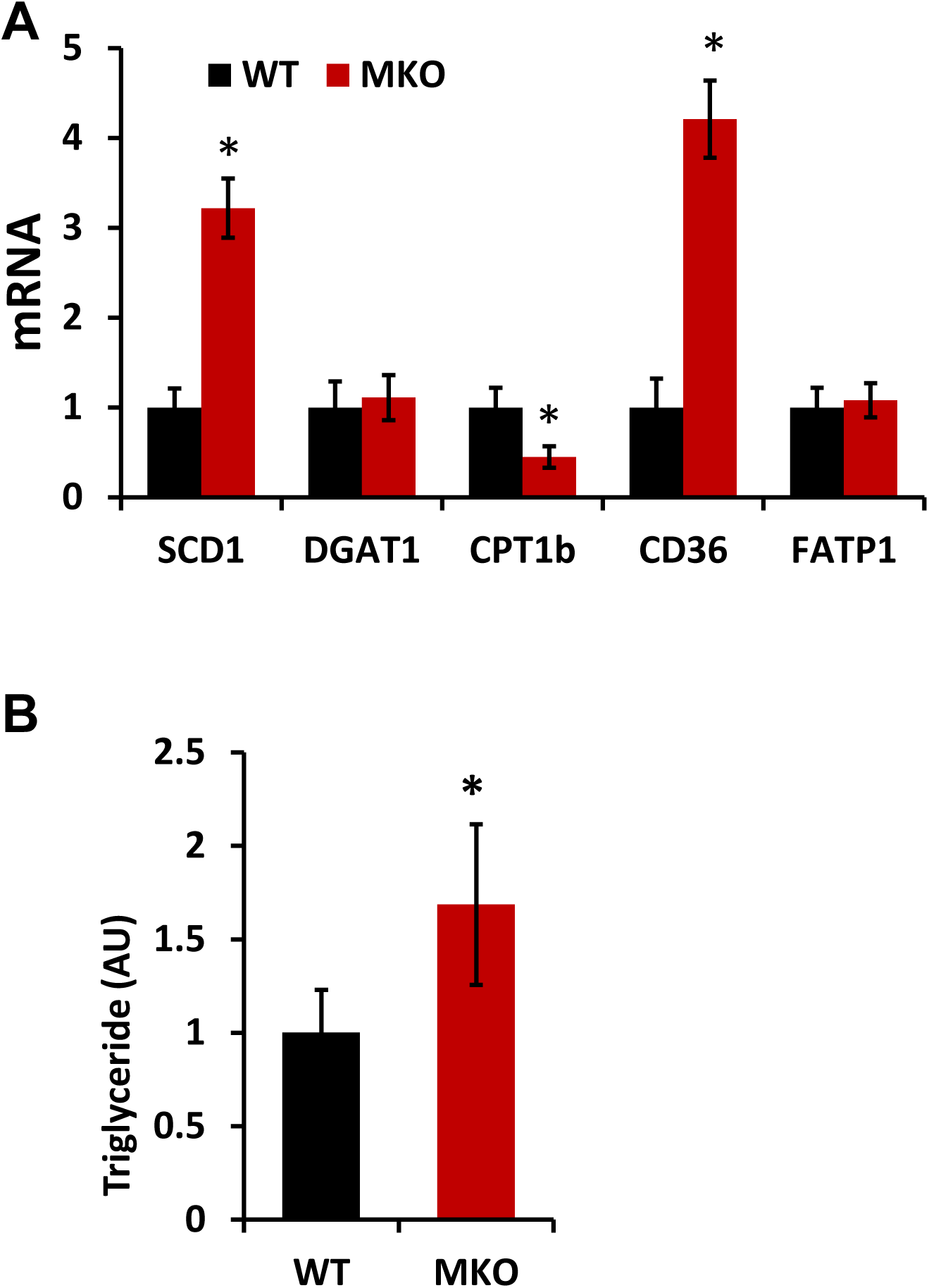
A loss of IPMK increases lipid accumulation. **A.** Gene expression in gastrocnemius by RT-qPCR (n=6). **B.** TG contents in the gastrocnemius muscle (n=8). Measurement was normalized by protein concentration. Data are mean +/- SEM, *p<0.05; Stearoyl-CoA desaturase-1; SCD1 (a rate limiting enzyme in the formation of monounsaturated FA). Diacylglycerol O-acyltransferase 1; DGAT1 (catalyzes the conversion of diacylglycerol and fatty acyl CoA to triacylglycerol). Carnitine palmitoyl transferase 1B; CPT1b (a rate liming enzyme for β-oxidation). Cluster of differentiation 36; CD36 (uptake of long chain fatty acids). Fatty acid transport protein 1; FATP1(uptake of long-chain fatty acids).

### Deletion of IPMK in skeletal muscle reduces exercise tolerance

Our data showed that deletion of IPMK interferes with glucose and lipid homeostasis and in particular, lipid utilization in the skeletal muscle. Thus, we proceeded to determine how the disrupted glucose and lipid metabolism in the skeletal muscle in MKO affects muscle function by conducting a forced treadmill exercise with 14-15 week-old WT and MKO mice before body weight significantly diverged. The treadmill test results showed that MKO mice had a significantly shorter running time until exhaustion than WT mice, suggesting that IPMK deficiency in muscle impaired their aerobic exercise capacity **(Fig. 4)**.

**Figure 4.**
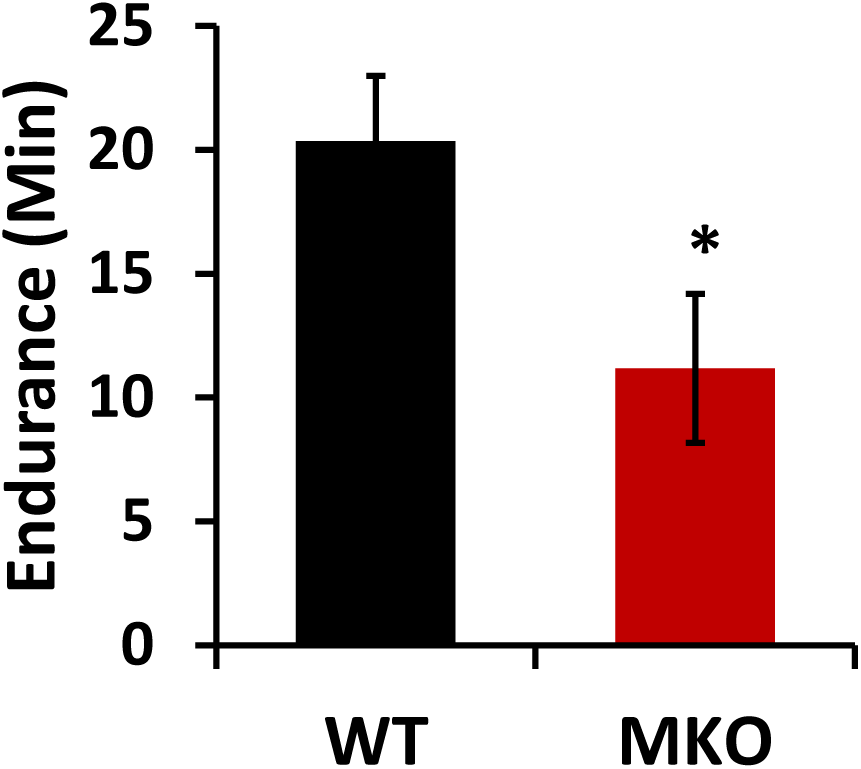
IPMK deletion in skeletal muscle reduces exercise capacity. WT and IPMK MKO male mice (14-15 week old) were subjected to exercise tolerance test. Data are mean +/- SEM, n=8. *p<0.05.

### A loss of IPMK impairs myocellular glucose utilization and insulin signaling

To better understand the cellular function of IPMK in muscle, we isolated primary myoblasts from *Ipmk*-loxp mice and differentiated them into myotubes after treating them with either Ad-GFP or Ad-Cre. First, we found that a loss of IPMK does not affect myocyte differentiation by checking the expression level of myogenin, a differentiation marker for myotube, between WT and *Ipmk^-/-^* cells. Interestingly, we found that the nuclear form of sterol regulatory element binding protein (nSREBP), a master regulator for lipid homeostasis, is up-regulated in *Ipmk^-/-^* cells while pAMPK is downregulated **(Fig. 5A).** Previously, we have shown that IPMK is necessary for glucose-mediated modulation of AMPK in various cell lines [7, 33]. Ubiquitously expressed AMPK senses the energy status of cells, regulates fuel availability, and plays a major in the regulation of glucose and lipids [27–30]. To further understand the role of IPMK in glucose homeostasis via AMPK in muscle, we acutely deleted IPMK using Ad-Cre in primary myocytes isolated from *Ipmk-*loxp. We found that a loss of IPMK attenuates glucose-mediated pAMPK inactivation in primary myocytes **(Fig.5B).** Moreover, glucose uptake, both basal and insulin-stimulated, was significantly decreased in *Ipmk^-/-^* myocytes compared to WT (**Fig. 5C**). Furthermore, we observed that insulin-mediated signaling events by the phosphorylation of Akt were inhibited in *Ipmk^-/-^* primary myocytes **(Fig. 5D).**

**Figure 5.**
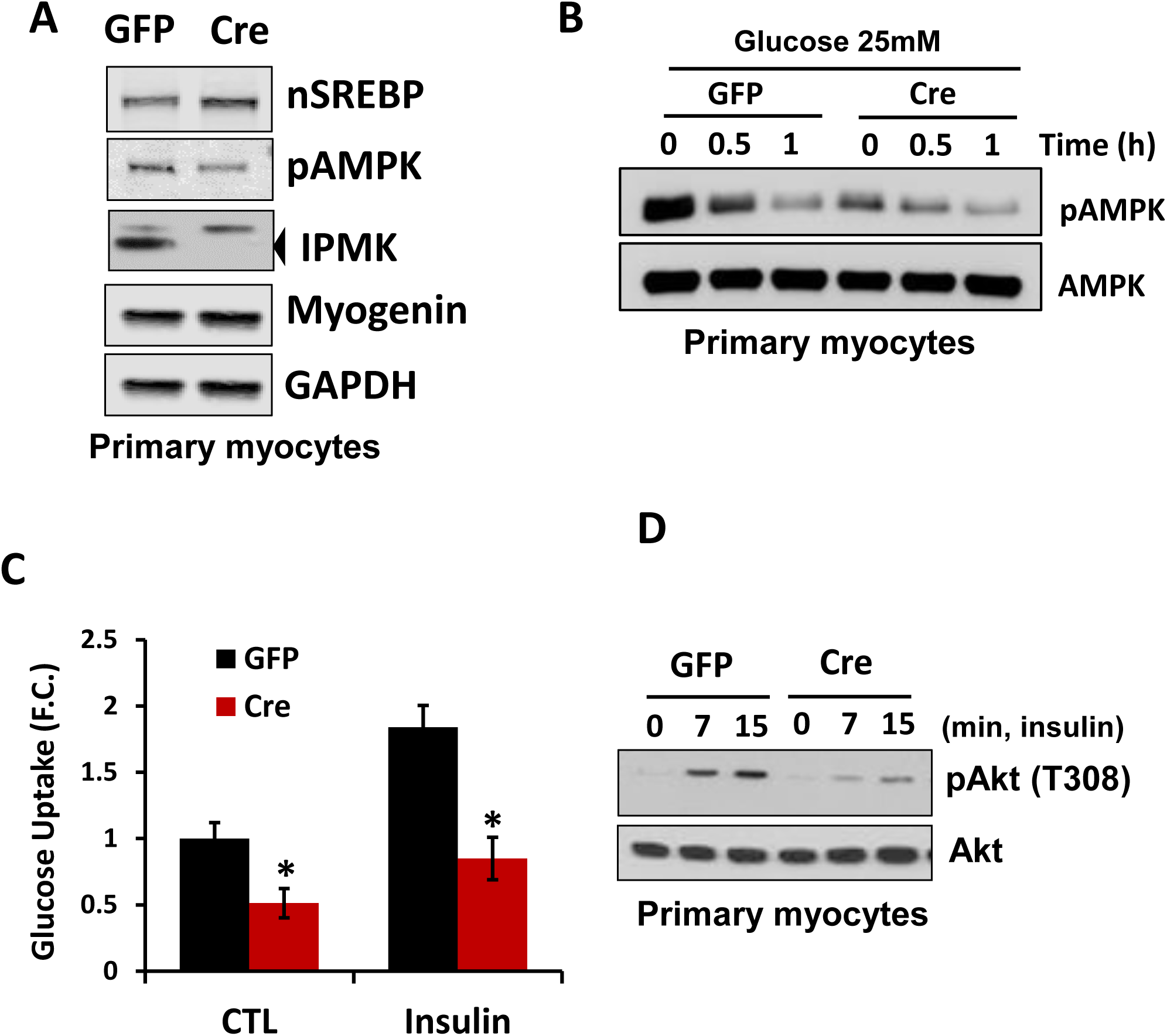
A loss of IPMK impairs myocellular glucose metabolism and insulin signaling. Primary myoblasts were isolated from *Ipmk*-loxp mice and treated with either Ad-GFP as a control or Ad-Cre. Then cells were differentiated into myotubes. **A**. Cell lysates were analyzed by Immunoblotting. **B.** Cells were pre-incubated in glucose-free buffer and incubated with 25 mM glucose for an indicated period of time. **C.** Cells were serum starved for 18 hrs and glucose uptake was measured in the presence of 100 nM insulin. **D.** Cells were serum starved for 18 hrs and were treated with 100 nM insulin for an indicated period of time. Data are mean +/- SEM, n=8. *p<0.05. Representative images from three independent trials for immunoblotting.

### A loss of IPMK disrupts lipid homeostasis in primary myocytes

We found that an acute loss of IPMK by Ad-Cre increased nSREBP in primary myocytes **(Fig. 5A).** To better understand whether IPMK plays a role in lipid metabolism, we examined levels of triglyceride (TG), fatty acid uptake, and fatty acid oxidation using primary myocytes with or without IPMK. First, we noted that the TG level is significantly increased in *Ipmk^-/-^* myocytes compared to WT. Moreover, there is an increase in ^14^C-oleic acid (OA) uptake and a decrease in β-oxidation, which was measured by ^14^CO_2_ released into media suggesting that a deletion of IPMK led to the disruption of lipid homeostasis in myocytes **(Fig.6)**.

**Figure 6.**
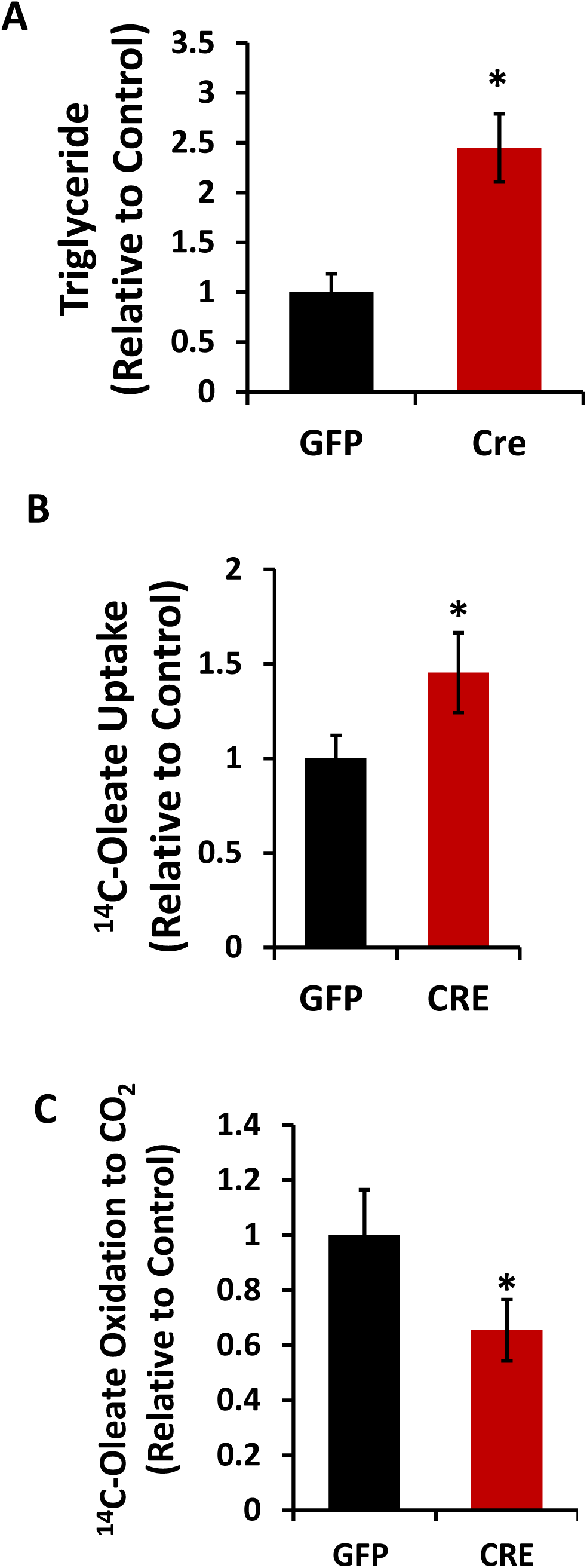
A loss of IPMK disrupts lipid homeostasis. Primary myoblasts were isolated from *Ipmk*-loxp mice and treated with either Ad-GFP as a control or Ad-Cre. Then cells were differentiated into myotubes. **A.** TG levels were measured. **B.** [1-^14^C]-oleic acid uptake (normalized by the protein concentration). **C.** β-oxidation of [1-^14^C]-Oleic acid was measured by CO_2_ released into media and data were normalized by the protein concentration. Data are mean +/- SEM, n=8. *p<0.05.

## Discussion

High levels of intramuscular triglyceride (IMTG) are common in obese or T2D individuals and are associated with insulin resistance [38, 39]. Additionally, it’s proposed that decreased fatty acid oxidation in obese individuals contributes to the accumulation of IMTG [40–42]. Consistent with these studies, we discovered that depletion of IPMK led to compromised insulin signaling, elevated fat accumulation, and reduced fat oxidation in myocytes. More importantly, skeletal muscle-specific IPMK-deficient mice exhibit reduced exercise capacity and increased body weight. The whole-body energy homeostasis analysis of IPMK-deficient mice suggests that weight gain in IPMK-deficient mice likely is linked to decreased EE. Abnormal fat accumulation and the reduction of endurance muscle function in IPMK-deficient muscle resemble insulin resistance-induced muscle weakness. Therefore, our study highlights the role of IPMK as a key regulator for the oxidative metabolism and exercise capacity of skeletal muscle.

Skeletal muscle is the primary tissue responsible for insulin-responsive glucose tolerance and lipid oxidation, making the maintenance of its functioning mass a crucial and dynamic process [31, 43]. Patients with T2DM and insulin resistance often exhibit reduced responsiveness to insulin stimulation of glucose uptake in skeletal muscle, poor exercise performance and increased fat accumulation [44–46]. Additionally, many studies have demonstrated that diet-induced or genetic insulin resistance specifically in the muscle of mice results in the development of glucose intolerance associated with elevated circulating triglycerides [47, 48]. Overexpression of kinase-dead insulin receptor (IR) or IGF-1 receptor (IGF1R) in muscle led to glucose intolerance, high levels of circulating triglyceride, and insulin resistance in mice [49, 50]. Our previous studies have shown that the deletion of IPMK reduced the activity of insulin signaling in hepatocytes and exacerbated high fat-induced insulin resistance in mice [35, 51]. Consistent with these findings, the depletion of IPMK in primary myocytes showed reduced AKT activation in response to insulin, and IPMK-MKO mice developed glucose intolerance, similar to findings in mice with overexpression of kinase-dead insulin receptor (IR) or IGF-1 receptor (IGF1R). Whole-body metabolism analysis of IPMK-MKO mice suggests that skeletal muscle depletion of IPMK results in a tendency for a decrease in total energy expenditure (Fig. 1), consistent with increased fat mass and unchanged locomotion (Fig. 1). Additionally, the respiratory exchange ratio in MKO mice exhibits significantly altered fatty acid utilization without any significant change in food intake (data not shown) during the light phase compared to WT mice. Taken together, these data indicate that IPMK deficiency in muscle alters insulin signaling, whole-body energy expenditure and fuel utilization.

Previously, we showed that IPMK interacts with AMPK [7, 33]. AMPK is a key sensor of cellular energy status and is activated in response to low energy levels, specifically when the AMP/ATP ratio increases [23, 27]. AMPK activation enhances lipid oxidation and reduces the incorporation of fatty acids into TG by phosphorylating ACC in skeletal muscle [23, 52, 53]. Conversely, the ablation of AMPK decreases fatty acid oxidation, increases lipid accumulation in skeletal muscle, and leads to elevated TG content [54]. Similarly, the reduction of AMPK activation in IPMK-deficient myocytes results in decreased glucose uptake (Fig. 5) and increased triglyceride accumulation. (Fig. 6), mirroring the phenotypes observed in AMPK-deficient muscles. Furthermore, IPMK-deficient myocytes exhibit increased expression of lipogenic genes, including CD36 and SCD1, while the expression of oxidative genes, such as CPT1b, is decreased. These results recapitulate the phenotypes of the AMPK-deficient mouse model, suggesting that the IPMK-AMPK axis may play a major role in the metabolic processes and functions of muscle.

IPMK deficiency leads to reduced oxidation and triglyceride accumulation in myocytes. Fat accumulation, along with insulin resistance, encourages the development of defects in fatty acid metabolism. These defects may include alterations in fatty acid oxidation, uptake, TG synthesis, or any combination of these processes in skeletal muscles. A reduction in oxidative enzyme activity and an increase in glycolytic activity in muscle cells indicate a shift from oxidative (aerobic) metabolism to glycolytic (anaerobic) metabolism, resulting in a decreased capacity for aerobic energy production. This shift can lead to reduced endurance [55–57]. The ability of IPMK to regulate the metabolic capacity in skeletal muscle may be achieved, at least in part, through genes of oxidative metabolism transcription in response to various environmental conditions like age or exercise. Given that decreased EE and exercise capacity were already observed in IPMK KO mice (Figs. 5 and 6), it is likely that altered muscle function could be attributed to the disrupted energy metabolism in IPMK-deficient mice. Moreover, it is prudent to point out that the observed reduction in endurance does not translate into heart dysfunction, as MLC-Cre specifically influenced skeletal muscle, not cardiac muscle [58].

In summary, our study demonstrates for the first time that depletion of IPMK in skeletal muscle disrupts glucose homeostasis and increases lipid accumulation with impaired insulin signaling. Additionally, our findings reveal that IPMK plays a significant role in regulating oxidative metabolism and exercise capacity in muscle. These results suggest a plausible mechanism for how muscle metabolic function may affect metabolic diseases and/or exercise capacity.

## Acknowledgment

We thank Jade West and Bouchra Tiab for their technical support. This work was supported by grants from the National Institutes of Health/National Institute of Diabetes and Digestive and Kidney Diseases (NIH/NIDDK) (SFK and RSA, DK135751).

## Notes

### Competing Interest Statement

The authors have declared no competing interest.

### Summary of Updates

This version of the manuscript has been revised to update some of the figures.

